# Creation and multi-omics characterization of a genomically hybrid strain in the nitrogen-fixing symbiotic bacterium *Sinorhizobium meliloti*

**DOI:** 10.1101/296483

**Authors:** Alice Checcucci, George C. diCenzo, Veronica Ghini, Marco Bazzicalupo, Anke Becker, Francesca Decorosi, Johannes Döhlemann, Camilla Fagorzi, Turlough M. Finan, Marco Fondi, Claudio Luchinat, Paola Turano, Tiziano Vignolini, Carlo Viti, Alessio Mengoni

## Abstract

Many bacteria, often associated with eukaryotic hosts and of relevance for biotechnological applications, harbour a multipartite genome composed by more than one replicon. Biotechnologically relevant phenotypes are often encoded by genes residing on the secondary replicons. A synthetic biology approach to developing enhanced strains for biotechnological purposes could therefore involve merging pieces or entire replicons from multiple strains into a single genome. Here we report the creation of a genomic hybrid strain in a model multipartite genome species, the plant-symbiotic bacterium *Sinorhizobium meliloti*. In particular, we moved the secondary replicon pSymA (accounting for nearly 20% of total genome content) from a donor *S. meliloti* strain to an acceptor strain. The *cis*-hybrid strain was screened for a panel of complex phenotypes (carbon/nitrogen utilization phenotypes, intra- and extra-cellular metabolomes, symbiosis, and various microbiological tests). Additionally, metabolic network reconstruction and constraint-based modelling were employed for *in silico* prediction of metabolic flux reorganization. Phenotypes of the *cis*-hybrid strain were in good agreement with those of both parental strains. Interestingly, the symbiotic phenotype showed a marked cultivar-specific improvement with the *cis*-hybrid strains compared to both parental strains. These results provide a proof-of-principle for the feasibility of genome-wide replicon-based remodelling of bacterial strains for improved biotechnological applications in precision agriculture.

## INTRODUCTION

Interest in large-scale genome modification and synthetic bacterial chromosome construction has strongly increased over the last decade (for instance see ^1^) with a goal of engineering bacterial strains with new or improved traits. However, phenotypes are often the result of the coordinated function of many genes acting together in a defined genome architecture ^2^. Hence, the ability to predict the phenotypic outcomes of large-scale genome modification requires a precise understanding of the genetic and regulatory interactions between each gene or gene product in the genome. As such, there is a need for integrated approaches, combining experimental evidences with computational-based methods, to interpret and potentially predict the outcomes of genome-wide DNA manipulations.

In this context, multipartite (or divided) genomes (i.e., genomes possessing more than one informational molecule) are particularly interesting. The genome of bacteria with a multipartite structure is typically composed of a principal chromosome that encodes the core housekeeping and metabolic genes essential for cellular life, and one (or more) secondary replicons (termed chromids and megaplasmids). More than 10% of the presently sequenced bacterial genomes are characterized by the presence of a multipartite architecture ^3, 4^. The secondary replicons can account for up to half of the total genome size, and their level of integration into cellular regulatory and metabolic networks is variable ^5, 6, 7{González, 2006 #3279^. In some cases, strong replicon-centric transcriptional networks have been suggested ^8, 9^. The apparently functional modularity of secondary replicons is particularly attractive from both ecological and biotechnological viewpoints. Indeed, secondary replicons might act as plug-and-play functional modules, potentially allowing the recipient strain to obtain previously untapped genetic information ^10^. This, in turn, might allow the emergence of novel phenotypic features leading, for example, to the colonization of a new ecological niche ^11^. Moreover, such modularity paves the way for large-scale, genome-wide manipulations of bacterial strains with multipartite genome structure, by synthetically merging complex biotechnologically important traits in the same strain ^12^.

However, it remains unclear to what extent complex phenotypes can be directly transferred into a recipient strain, as secondary replicons are in part co-adapted to the host genome, for example, through regulatory interaction and/or inter-replicon metabolic cross-talk ^8, 13^. We are aware of only one study examining the phenotypic consequences of replacing a large (> 800 kb) native secondary replicon with a homologous replicon of closely related strains or species. In that study, the third replicon of *Burkholderia cepacia complex* strains was mobilized and the effects on various phenotypes including virulence was examined ^14^. It was found that in some cases, phenotypes were dependent solely on the secondary replicon, whereas in other cases, the phenotypes depended on genetic/regulatory interactions with the other replicons ^14^. However, additional studies are required to examine the generalizability of those observations.

To further test the feasibility, the stability, and the predictability of secondary replicon shuffling on the phenotype(s) of the cell, here we have performed experimental and *in silico* replicon transplantation between two bacterial strains. We used the symbiotic nitrogen-fixing bacterium *Sinorhizobium meliloti* as the model, given that it has a well-studied multi-replicon genome structure ^11, 15^. Additionally, *S. meliloti* represents a highly valuable microorganism in agriculture, as its symbiosis with crops like alfalfa is estimated to be worth more than $70 million/year in the U.S.A. ^16^. The genome of the mostly commonly studied *S*. *meliloti* reference strains (Rm1021 and Rm2011) is composed by a chromosome (~ 3.7 Mb), and two secondary replicons: a chromid (~ 1.7 Mb, called pSymB, carrying several genes involved in rhizosphere colonization) and a megaplasmid (~ 1.4 Mb, called pSymA, carrying most of classical symbiosis genes). *S. meliloti* large replicons have recently been proposed as scaffolds for novel shuttle vectors for synthetic biology ^17^. Furthermore, genome reduction experiments previously performed have led to the complete removal of one or both of the two secondary replicons^11, 18^, and an *in silico* genome-scale metabolic model has been reconstructed ^19^, paving the way for massive genome-scale remodelling of *S. meliloti*.

Here, we constructed a hybrid strain containing the chromosome and the chromid of the laboratory *S. meliloti* Rm2011 strain with the pSymA replicon from the wild isolate *S. meliloti* BL225C. The genome of BL225C is 290 kbps larger than that of Rm2011 (which has a genome highly similar to Rm1021 strain) ^15, 20^, and 1,583 genes are present in only one of these strains^21^are present in only one of these strains ^22^. Furthermore, the BL225C strain has been shown to have several interesting biotechnological features, including plant growth promotion, and nodulation efficiency ^15, 23^. The majority of the genetic differences between Rm2011 and BL225C strains is associated with the symbiotic pSymA homolog megaplasmids; 836 of the 1,583 variable genes are located on this replicon^21^. We can then expect that creating a hybrid strain between Rm2011 and BL225C, by moving the pSymA-equivalent from BL225C to the Rm2011 derivative lacking pSymA, will provide a good testing ground for i) feasibility of large replicon shuffling between strains, and ii) stability and predictability of the phenotypes linked to such replicons. We term this novel hybrid strain as *cis-*hybrid since it derives from *cis*-genic manipulation, indeed it contains genetic material from the pangenome pool of the same species (in contrast to a *trans*-genic strain that would contain genes from a distinct species). *Cis*-hybrid strains could be an important way to promote environmental-friendly and regulatory compliant biotechnology and synthetic biology in bacterial species of interest in agricultural and environmental microbiology ^12^.

## RESULTS AND DISCUSSION

We report here the creation of a *cis*-hybrid *S. meliloti* strain, where the symbiotic-related megaplasmid pSINMEB01, homologous to pSymA (~ 1.6 Mb in size, accounting for nearly 23% of total genome) was transferred from the natural strain BL225C to the laboratory strain Rm2011. *In silico* metabolic reconstruction and a large set of phenotypic tests, including Nuclear Magnetic Resonance (NMR)-based metabolomic profiling, Phenotype Microarray™, and symbiotic assays with different host plant cultivars have been performed, as described in the following paragraphs.

### Experimental creation of a *cis*-hybrid strain

Starting with the derivative of the *S. meliloti* Rm2011 strain that lacks pSymA replicon, herein referred to as ∆pSymA ^24^, we produced a *cis*-hybrid strain that contains the Rm2011 chromosome and pSymB chromid, and the pSymA replicon from a genetically and phenotypically distinct *S. meliloti* strain, BL225C (all strains used in this work are listed in Table 1)^15, 23^. The *cis*-hybrid strain was produced through a series of conjugations as described in the Methods section. Briefly, a plasmid for over-expressing *rctB* ^25^was transferred to BL225C; as RctB is a negative regulator of RctA ^25^, which in turn is a negative regulator of the pSymA conjugal genes ^26^, this step was necessary to promote pSymA transfer without mutating the replicon. Concurrently, a plasmid carrying an antibiotic resistance gene marker (gentamicin, plasmid pMP7605) was transferred to the ∆pSymA strain to allow for the use of the gentamicin resistance marker in the selection of *cis*-hybrid transconjugants in the next step. Finally, a mating mixture of the Rm2011 acceptor strain and the BL225C donor strain was prepared, and *cis*-hybrid transconjugants were isolated on a medium selective for the gain of the pSymA replicon (see Methods). Correct construction of the *cis*-hybrid strains was initially confirmed through PCR amplification on specific unique genes on the Rm2011 chromosome and pSymB, and the pSymA homolog of BL225C (pSINMEB01) (see Table S1). Subsequently, whole genome sequencing (Figure 1) confirmed the complete transfer of pSymA, and the banding pattern observed in pulse-field gel electrophoresis (PFGE) (Supplemental Figure S1) was consistent with pSINMEB01 being present as an independent replicon (i.e., not integrated into the chromosome or pSymB).

**Table 1.**
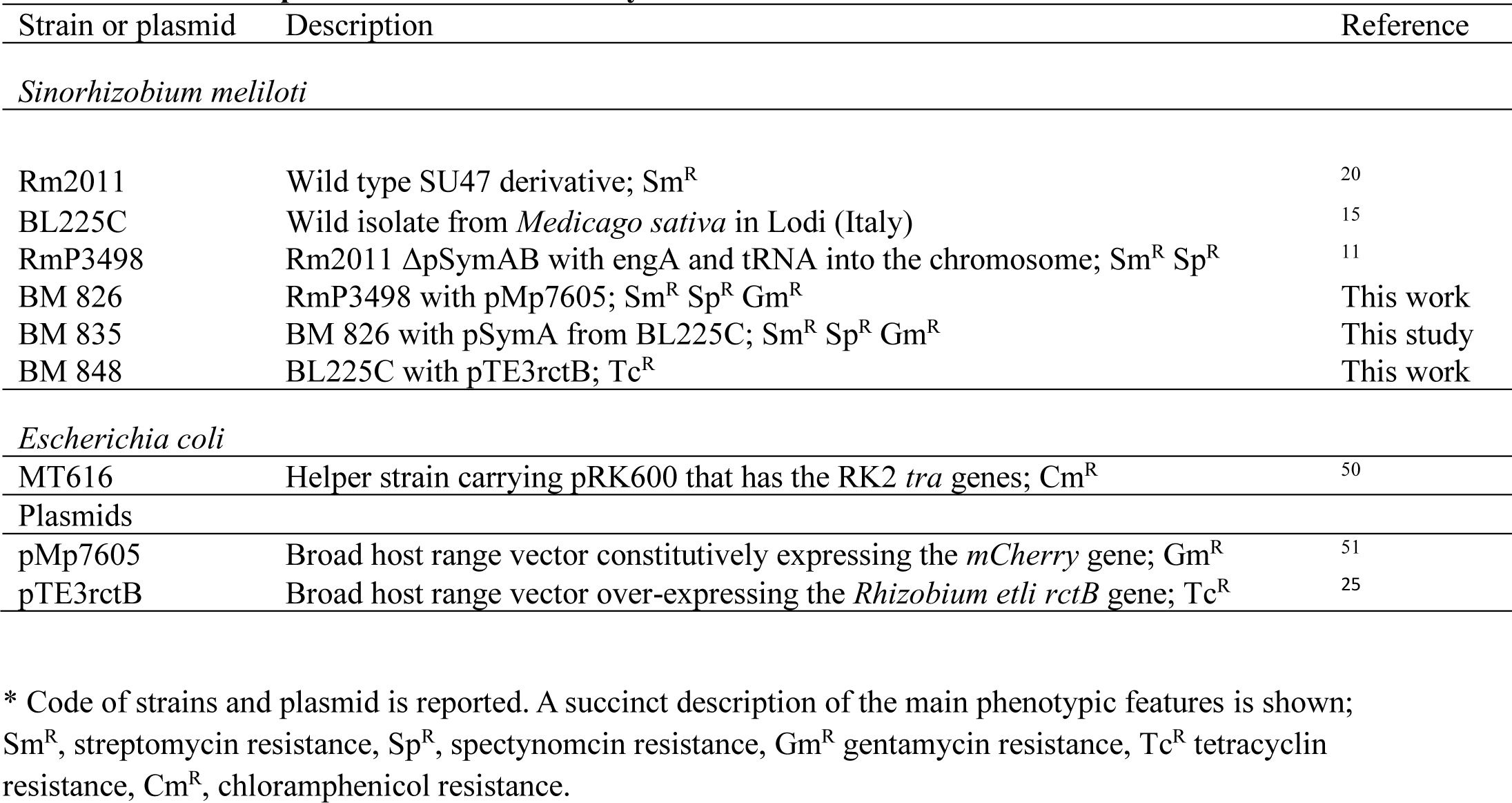
**Strains and plasmids used in this study\***

| Strain or plasmid | Description | Reference |
| --- | --- | --- |
| *Sinorhizobium meliloti* |  |  |
| Rm2011 | Wild type SU47 derivative; Sm<sup>R</sup> | 20 |
| BL225C | Wild isolate from *Medicago sativa* in Lodi (Italy) | 15 |
| RmP3498 | Rm2011 ΔpSymAB with engA and tRNA into the chromosome; Sm<sup>R</sup> Sp<sup>R</sup> | 11 |
| BM 826 | RmP3498 with pMp7605; Sm<sup>R</sup> Sp<sup>R</sup> Gm<sup>R</sup> | This work |
| BM 835 | BM 826 with pSymA from BL225C; Sm<sup>R</sup> Sp<sup>R</sup> Gm<sup>R</sup> | This study |
| BM 848 | BL225C with pTE3rctB; Tc<sup>R</sup> | This work |
| *Escherichia coli* |  |  |
| MT616 | Helper strain carrying pRK600 that has the RK2 *tra* genes; Cm<sup>R</sup> | 50 |
| Plasmids |  |  |
| pMp7605 | Broad host range vector constitutively expressing the *mCherry* gene; Gm<sup>R</sup> | 51 |
| pTE3rctB | Broad host range vector over-expressing the *Rhizobium etli rctB* gene; Tc<sup>R</sup> | 25 |

\* Code of strains and plasmid is reported. A succinct description of the main phenotypic features is shown; Sm<sup>R</sup>, streptomycin resistance, Sp<sup>R</sup>, spectinomycin resistance, Gm<sup>R</sup> gentamycin resistance, Tc<sup>R</sup> tetracyclin resistance, Cm<sup>R</sup>, chloramphenicol resistance.

**Figure 1.**
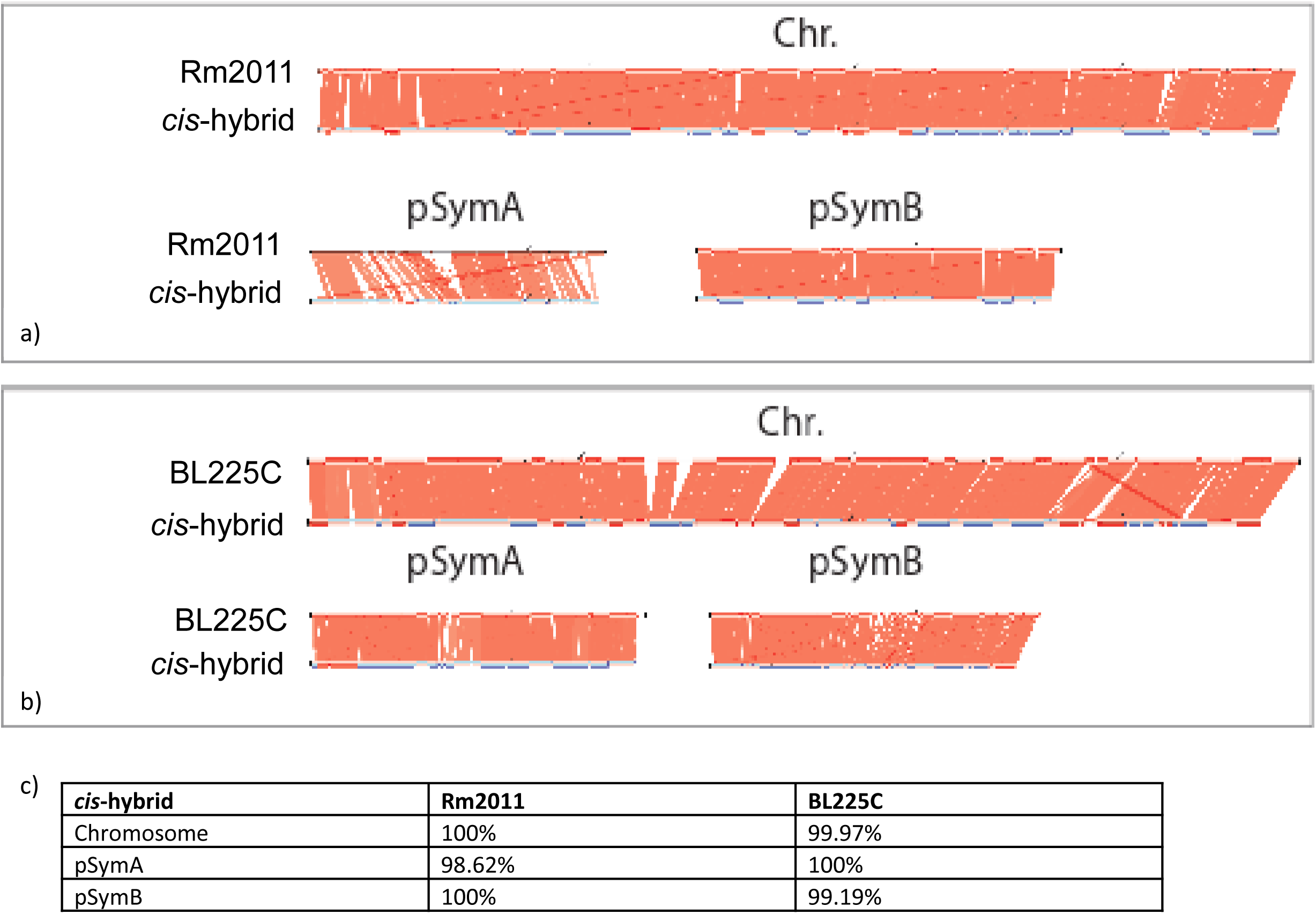
(a) Comparison of *cis*-hybrid strain genome sequence with 2011 chromosome and pSymB, and with pSymA of the donor strain (BL225C) (contigs alignment performed with Contiguator (Galardini*, et al.*, 2011); (b) Percentage of identity of each replicon which composed the multipartite genome of *cis*-hybrid strain with those of the donor strains (Rm2011 and BL225C).

### *In silico* metabolic network reconstruction

In addition to the experimental creation of the *cis-*hybrid strain, we attempted to predict the metabolic outcomes of the *cis-*hybrid strain by generating a new metabolic model which includes the genomic features present in the *cis-*hybrid strain. The curated iGD1575 reconstruction (herein referred to as the Rm2011 reconstruction) was used to represent metabolism of *S. meliloti* Rm2011 ^24^; although iGD1575 is based on the strain Rm1021, the genomic content of these strains are 99,9% identical, with the exception of numerous SNPs ^20^ that are not considered in the process of metabolic reconstruction. Next, our recently described pipeline ^27^ was used to build a representation of BL225C based on a draft reconstruction built with the Kbase webserver and enhanced based on the iGD1575 model. An *in silico* representation of the *cis*-hybrid strain was then built by removing all pSymA genes (and dependent reactions) from the Rm2011 model, followed by the addition of all pSymA (pSINMEB01) genes (and associated reactions) from the BL225C model using our published pipeline ^27^. Despite there being numerous (47 to 143 gene) differences in the gene contents of the metabolic reconstructions, the Rm2011 model differed from the BL225C and the cis-hybrid models by no more than a half dozen reactions (Table 2). The low reaction variability between models may i) reflect the difficulty in predicting the function of the *S. meliloti* variable gene content, ii) suggest the presence of non-orthologous genes encoding proteins catalyzing the same reaction(s), and/or iii) indicate that few metabolic features are dependent on the accessory gene set. Not surprisingly, given the near identical reaction content of the reconstructions, the outputs of flux balance analysis simulations for the different reconstructions were nearly identical (data not shown); therefore, we do not describe these results further.

**Table 2.** **Comparison of *S. meliloti* metabolic network reconstructions.** The gene and reaction content of the four *S. meliloti* metabolic reconstructions used in this work are shown. For each cell, the values are a comparison of the strain indicated on the left with the strain indicated along the top. Three values are provided in each cell, and these correspond to the following. The first value is the number of genes or reactions in common between the models. The second value is the number of genes or reactions present in the reconstruction on the left but not in the one along the top. The third value is the number of genes or reactions present in the reconstruction along the top but not in the one on the left.

| Strain | Genes |  |  | Reactions |  |  |
| --- | --- | --- | --- | --- | --- | --- |
| | BL225C | cis-hybrid | ΔpSymA | BL225C | cis-hybrid | ΔpSymA |
| Rm2011 | 1525 / 52 / 91 | 1551 / 26 / 76 | 1336 / 241 / 0 | 1821 / 6 / 6 | 1823 / 4 / 6 | 1755 / 72 / 0 |
| BL225C | - | 1598 / 18 / 29 | 1308 / 308 / 28 | - | 1827 / 0 / 2 | 1753 / 74 / 2 |
| cis-hybrid | - | - | 1336 / 291 / 0 | - | - | 1755 / 74 / 0 |

### Metabolic phenotypes and profiles of the *cis*-hybrid strain

Phenotype MicroArray™ experiments were performed to test the growth of the *cis*-hybrid strain, as well as the parental and wild type strains, with 192 different carbon sources and 95 different nitrogen sources. Previous work has shown that the pSymA megaplasmid has little contribution to the metabolic capacity of *S. meliloti* ^11,18^. Consistent with this, only minor changes in the metabolic growth abilities were observed following the introduction of the pSymA of BL225C (pSINMEB01) into the ∆pSymA strain (Figure 2/Table S2). This confirms that transplantation of pSymA did not result in a major disturbance in the metabolic abilities of the recipient strain. Additionally, the ∆pSymA strain lost the ability to use 3-methylglucose as a carbon source and cytosine as a nitrogen source, and both abilities were restored upon the introduction of the pSymA replicon from BL225C. This result helps validate the replicon transplantation approach by confirming that at least some of the genes on the pSymA replicon of BL225C are properly expressed, and their gene productions functional, in the Rm2011 background.

**Figure 2.**
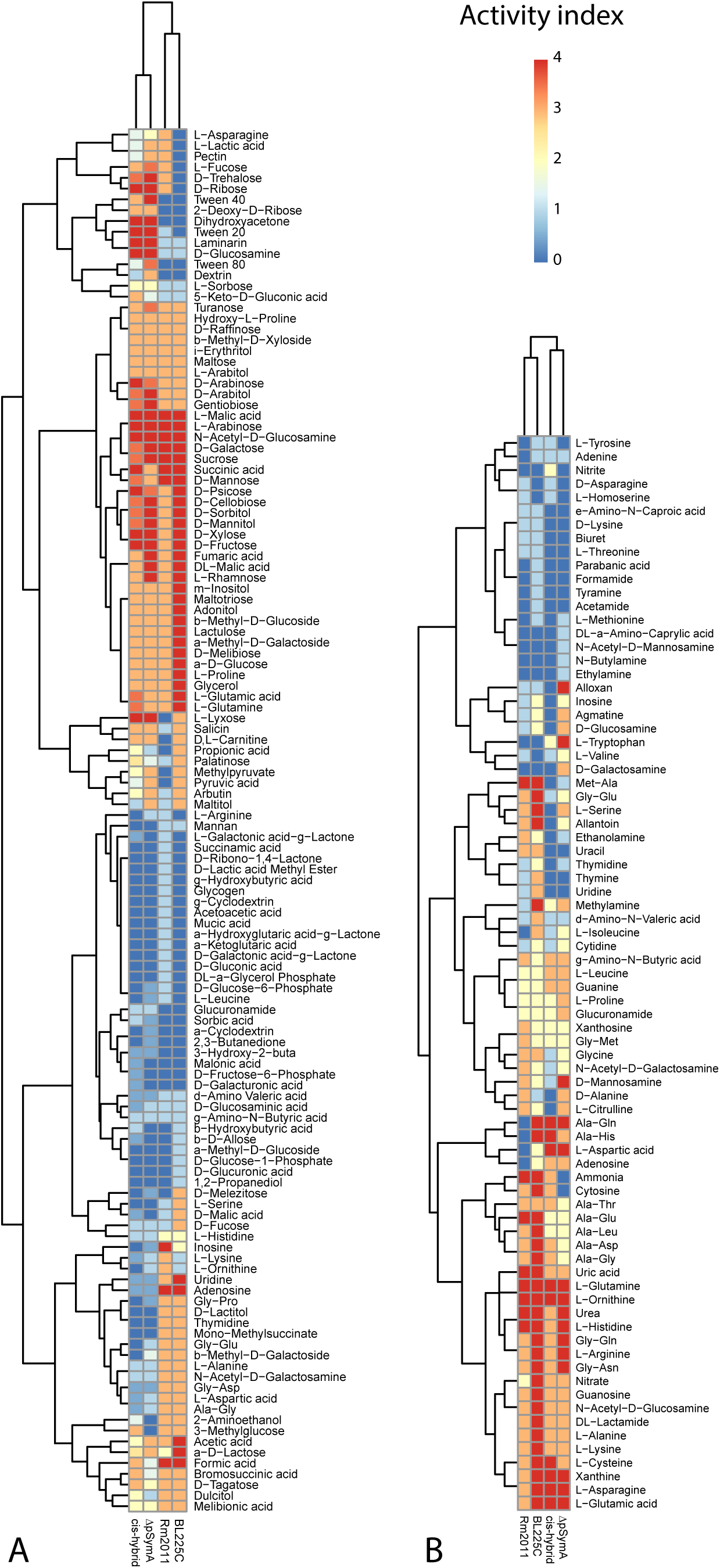
Metabolic phenotype of the *cis*-hybrid strain. Heatmap of Phenotype Microarray profiles of the growth on different carbon and nitrogen sources for Rm2011, BL225C, *cis*-hybrid and ΔpSymA strains. (a) heatmap with Euclidean clustering; (b)values of pairwise Euclidean distances

A metabolomic analysis through ^1^H nuclear magnetic resonance (NMR) was performed to further define the metabolic consequences of pSymA transplantation. Using an untargeted approach, both cellular lysates and spent growth media were analysed to identify the fingerprint of the endo- and exo-metabolomes of the two parental strains, the *cis*-hybrid strain, and the ∆pSymA recipient.

PCA was used to generate an initial overview of the metabolome differences among the four strains (Figures 3.a and 3.b), followed by PCA-CA to obtain the best discrimination among the strains by maximizing the differences among their metabolomic profiles (Figures 3.c and 3.d). In both the PCA and PCA-CA score plots (Figure 3), the c*is*-hybrid strain clustered very close to both the ∆pSymA recipient strain and to the parental strain Rm2011, whereas the parental donor strain BL225C clustered separately. These results are consistent with previous data indicating that pSymA has little contribution to the metabolome^6^, proteome^7^, or transcriptome ^28^ of *S. meliloti* Rm2011 in laboratory conditions. Importantly, these results confirmed that the synthetic large-scale horizontal gene transfer performed here to produce the *cis*-hybrid strain did not result in a major perturbation of the cellular metabolism.

**Figure 3.**
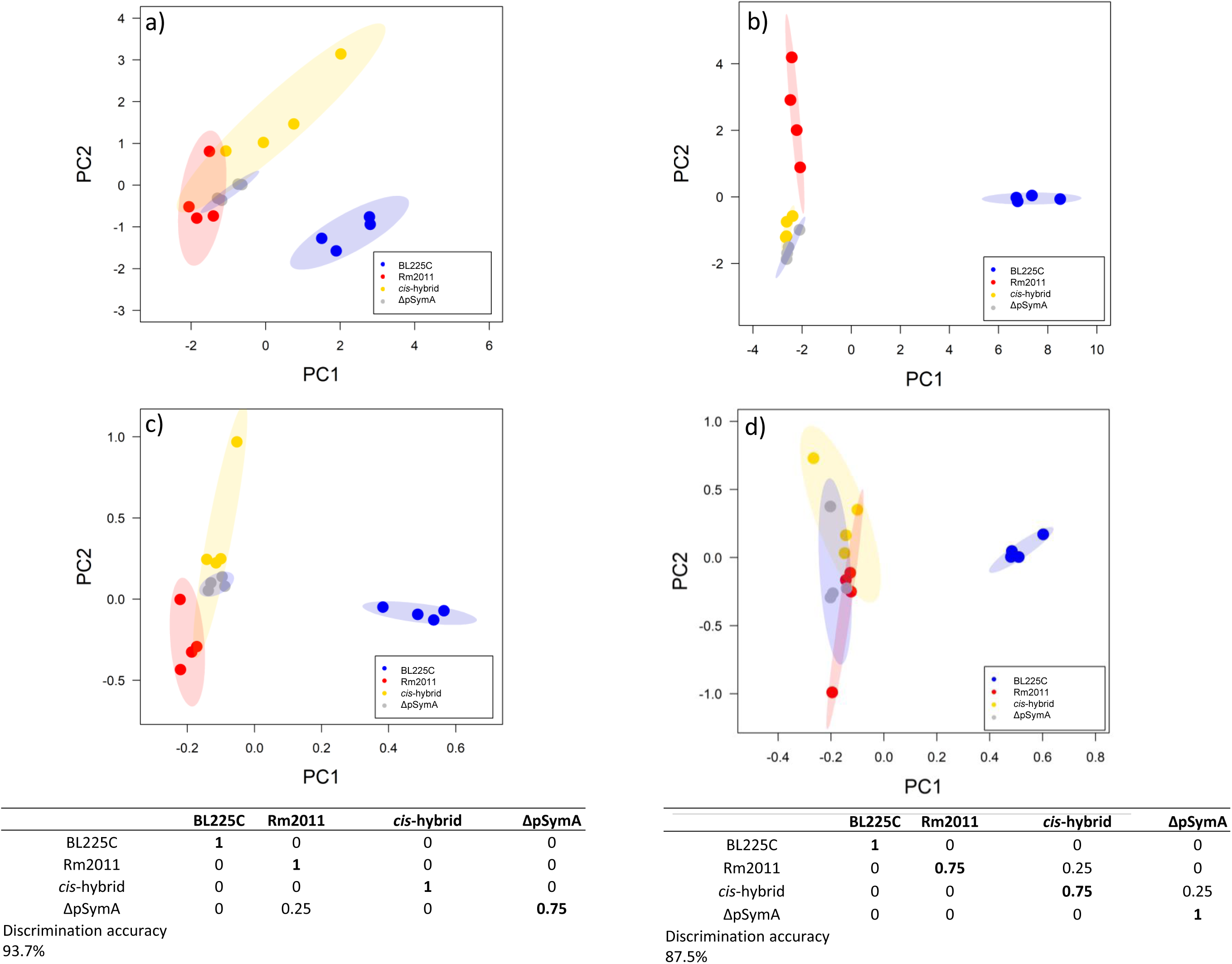
^1^H Nuclear Magnetic Resonance (NMR)-based metabolomic profiles of cellular lysates and growth media of Rm2011, BL225C, *cis*-hybrid and ΔpSymA strains. Score plot of PCA (a;b) and PCA-CA (c;d) analysis of cell lysates (a;c) growing media (b;d). The confusion matrices and the discrimination accuracy values for PCA-CA analysis are also reported. Ellipses in the score plots illustrate the 95% confidence level.

In addition to the multivariate analysis of the metabolic NMR fingerprints described above, the signals of 25 and 19 metabolites were unambiguously assigned and integrated in the ^1^H-NMR spectra of the cell lysates and growth media, respectively (Figure S2). The metabolites that are characterized by statistically significant differences in concentration levels in at least one strain with respect to the two other strains are indicated in Supplementary Table S3 and are also reported in Supplemental Figure S3. Validating the ability of this approach to detect metabolic differences between the strains, it was noted that the ∆pSymA strain exported cytosine unlike the wild type Rm2011 or the *cis*-hybrid strain, consistent with the inability of this strain to catabolize cytosine as shown by the Phenotype MicroArray™ data.

### Assessment of the phenotypes of the *cis*-hybrid strain

#### Growth profiles in synthetic laboratory media

Growth profiles of the *cis*-hybrid and parental strains in complex (TY) and defined (M9-succinate) media are reported in Figure 4. In the complex TY medium (Figure 4.a), growth of the *cis*-hybrid strain was impaired relative to the recipient (∆pSymA), and to the Rm2011 and BL225C parental strains. Or in other words, gain of the BL225C pSymA replicon by the ∆pSymA strain resulted in a decrease in the growth rate in TY medium. Although we cannot provide a definitive explanation for this phenomenon, it may be that the simultaneous gain of hundreds of new genes not integrated into cellular networks imposes a high metabolic cost to the cell, resulting in impaired fitness. In contrast, little to no difference was observed in the growth rate of any of the strains in the defined M9-succinate medium (Figure 4.b). The lack of an observable growth impairment of the *cis*-hybrid strain in the M9-succinate medium may be due to the growth impairment being masked by the general decrease in growth rate of all strains in this medium. Moreover, the similarity of the growth profiles of all strains in the minimal medium suggest that, at least in artificial laboratory conditions, the primary growth characteristics of these strains are primarily dependent on the core, not accessory, genome.

**Figure 4.**
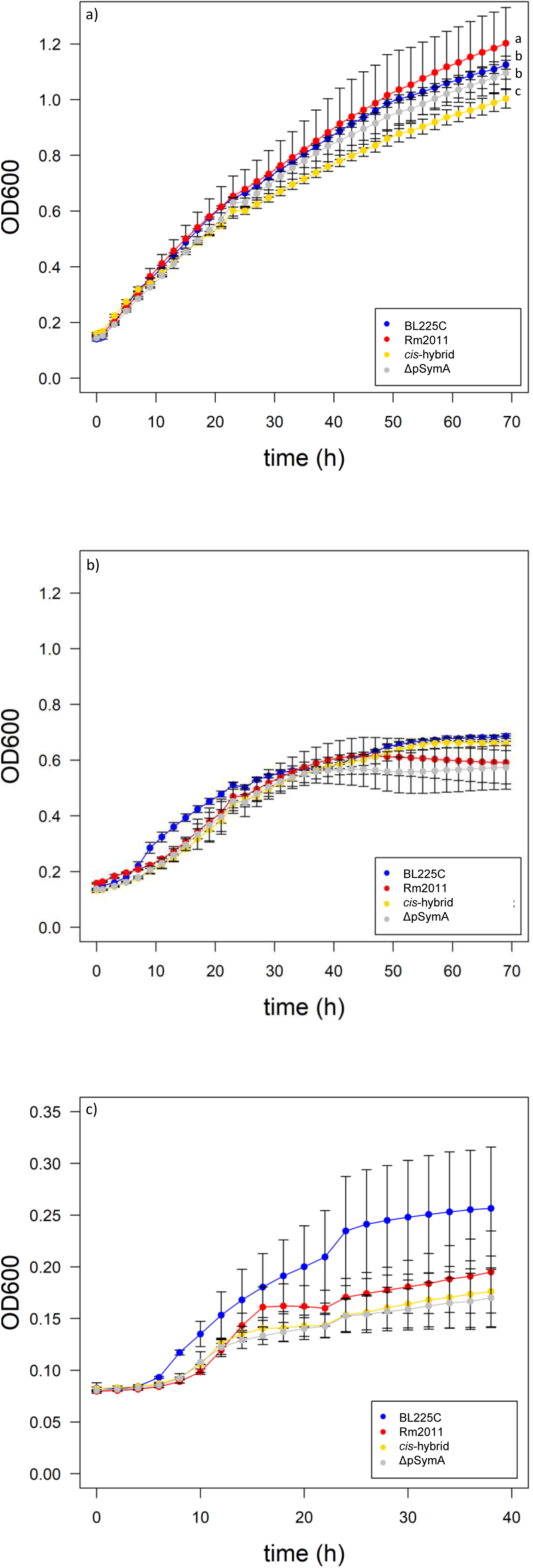
Growth phenotypes of the *cis*-hybrid strain. The growth of *S. meliloti* was examined in TY complex medium (a) M9 minimal medium (b) and (c) M9 +succinate and root exudates as sole N source. Data points represent averages from quadruplicate measurements. The letters on the curves represent the statistically significant differences among the strains growth (p<0.005, Tukey post-hoc contrasts).

#### Growth using root exudates as a nutrient source

Root exudates can be considered a proxy of the nutritional conditions of the plant rhizosphere ^29^. We therefore evaluated the ability of the three strains to grow on M9 mineral medium supplemented with root exudates of *Medicago sativa*, a *S. meliloti* symbiotic partner. None of the strains were able to grow when the root exudate was used as the sole carbon source; this was likely due to the root exudate being too dilute for use as a carbon source. In contrast, all strains could utilize the root exudate as the sole nitrogen source when provided succinate as a carbon source, and differential growth patterns were observed (Figure 4.c). In particular, BL225C displayed the highest growth among all four strains when grown with root exudates as a sole nitrogen source. Plating for viable colony forming units confirmed the differences in the final population densities (data not shown). As the robust growth of BL225C with root exudates did not transfer to the *cis*-hybrid strain, it is likely that this phenotype is primarily dependent on the chromosome and/or pSymB of BL225C, as was suggested by previous studies ^11, 21, 24, 30^. This observation would further suggest that the adaptation of the tested strains to growth in the rhizosphere occurred prior to the gain of pSymA and symbiotic abilities, consistent with recent work indicating that the majority of *S. meliloti* rhizosphere growth-promoting genes are chromosomally encoded ^31^. Finally, considering that there are relatively few differences in the nitrogen metabolic capacity of Rm2011 and BL225C ^23^, and that FBA (flux balance analysis) simulations for the metabolic model reconstructions were nearly identical (data not shown), we hypothesize that the growth differences observed between these strains is primarily related to regulatory differences, and less so to differences in metabolic genes.

#### Biofilm formation

Biofilm formation is a key factor in root colonization and plant invasion for many Proteobacteria^32^. In the light of producing a novel strain with good biotechnological features, biofilm formation was measured for the *cis*-hybrid strain. Biofilm production (estimated as the total biofilm-to-biomass ratios) by the *cis*-hybrid and the parental strains was similar (Figure S4). Interestingly, the ∆pSymA recipient strain showed a higher (p< 0.005) level of biofilm production compared to the other three strains. This led us to hypothesize that at least under the tested conditions, there is a pSymA mediated negative regulation of biofilm formation in both Rm2011 and BL225C strains. In contrast to these results, previous studies have suggested that deletion of pSymA ^33^, or just the pSymA-encoded common *nod* genes ^34^, results in a major reduction of biofilm formation. Future work is required to understand why the biofilm production phenotypes of *S. meliloti* strains lacking pSymA differed so dramatically between this study and those by Fujishige *et al*. ^42^.

### Symbiotic phenotypes of the *cis*-hybrid strain

Many of the key genes required for symbiotic abilities (e.g. nodule formation and nitrogen-fixation) are present on pSymA of *S. meliloti* Rm2011^35^ and the homologous megaplasmid pSINMEB01 of BL225C ^15^. These replicons additionally contain non-essential genes that promote improved symbiotic abilities^36^. While many of the symbiotic genes are conserved between these strains, relevant differences between pSymA and pSINMEB01 are present. The 482 genes exclusive to pSINMEB01 included symbiotic (e.g. *nws, hemA* homolog, *C P450 ^15^*) and nonsymbiotic functions (e.g. *acdS*) ^37^. For these reasons, this replicon swapping study was initiated in large part to evaluate whether swapping the symbiotic megaplasmid could promote differential symbiotic abilities.

To test the robustness of symbiotic abilities following replicon transplantation, *in vitro* symbiotic assays were performed on a panel of alfalfa cultivars, as alfalfa is the main host legume of *S. meliloti* ^38^(Figure 5). In particular, the *cis*-hybrid strain and the two parental strains (Rm2011, BL225C) were tested in combination with seven alfalfa cultivars (Table S4). These cultivars belong to the species *M. sativa*, *Medicago x varia*, and *Medicago. falcata*, and they are representative of the variability of cultivars and germplasm mainly used as crops in Europe. Moreover, BL225C was originally isolated on the *M. sativa* cultivar “Lodi” at the CREA-FLC institute (Italy) during a long-course experiment ^39^. The percentage of nodulated plants (Figure 5.a), the number of nodules per plant (Figure 5.b), the shoot dry weight (Figure 5.c), the plant aerial part length (Figure 5.d), and nodule colonization abilities (Supplemental Figure S5) were recorded using standard procedures ^23, 40^. Not surprisingly, for each strain there was high variability in the symbiotic phenotypes observed with the different cultivars. The symbiotic interaction is a multistep developmental process which involves a tight exchange of signals between the bacterium and the plant root at both rhizospheric and endophytic levels^12, 41^. Earlier works have demonstrated strain and cultivar specificities in this process, which result in *S. meliloti* strains displaying differential symbiotic effectiveness with various plant genotypes ^39, 42^.

**Figure 5.**
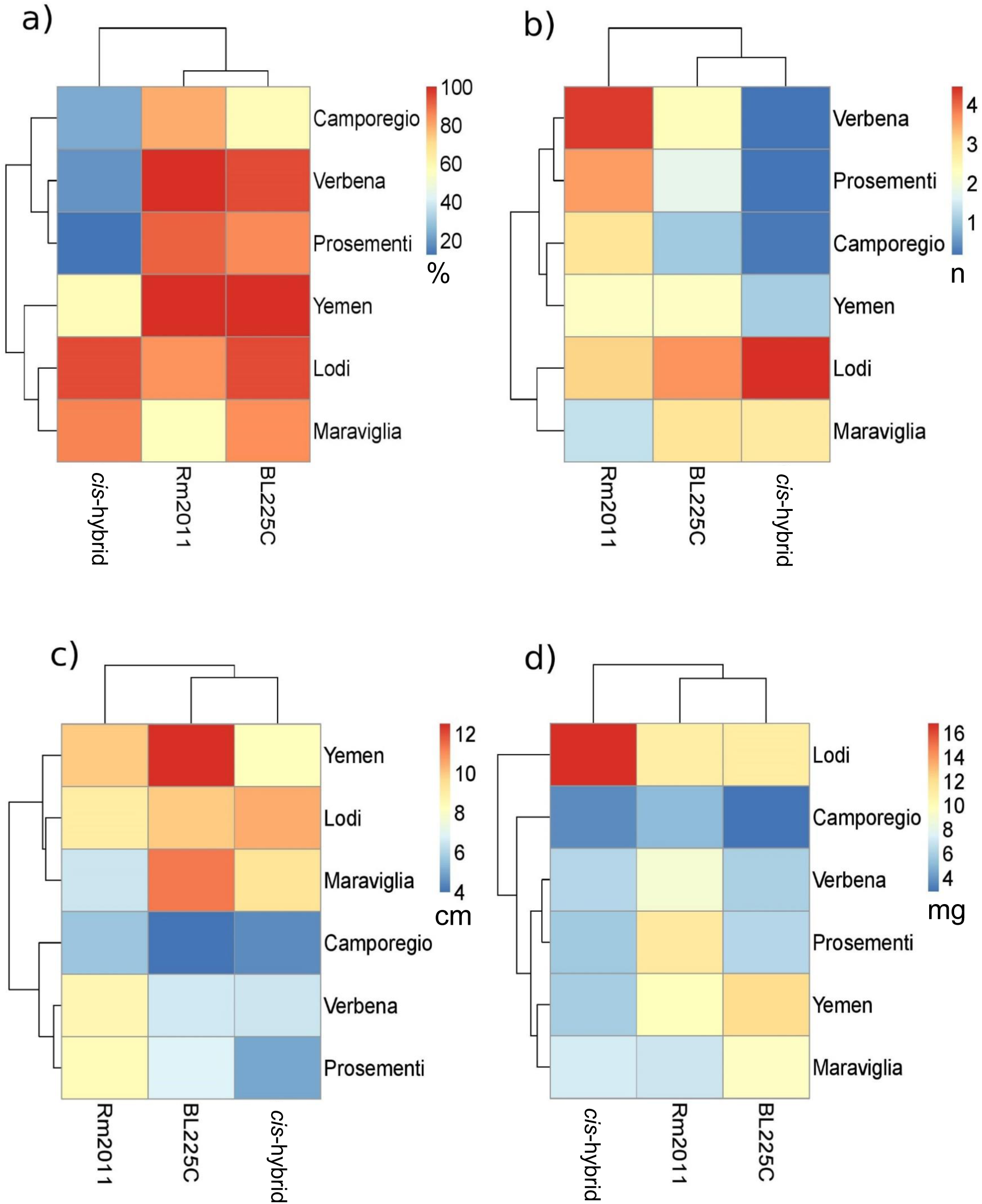
Symbiotic capabilities of the *cis*-hybrid strain. Heatmaps of symbiotic performances profiles for Rm2011, BL225C and cis-hybrid strains in a panel of seven alfalfa cultivars; (a) percentage of nodulated plants, (b) number of nodules per plant, (c) plant aerial part length (cm) and (d) the shoot dry weight (mg).

The *cis*-hybrid strain performed very poorly in symbiosis with some cultivars, such as in the cv. “Prosementi”, “Camporegio” and “Verbena”, in particular in the number of nodules per plant and the length of the aerial part (Figure 5b;c, Table S5) (p<0.005), indicating that the pSymA and pSINMEB01 megaplasmids are not always interchangeable. This could reflect the importance of the genomic context of the symbiotic megaplasmid and hypothetically the importance of inter-replicon regulatory networks ^8, 43^. Alternatively, pSINMEB01 may lack important (but still unknown) symbiotic genes with their function replaced by chromosomal genes in BL225C but not by chromosomal genes in Rm2011.

Strikingly, the *cis*-hybrid strain displayed clearly improved symbiotic capabilities during symbiosis with the cultivar “Lodi” compared to both Rm2011 and BL225C (Figure 5). This was true for several key measures of symbiosis, including nodule number per plant, shoot dry weight, and length of the aerial part of the plant. These data suggest the presence of nonlinear and genomic context dependent genic interactions in the establishment of symbiotic abilities. Such interactions may resemble (at the logic level) those present in some eukaryotic genomes that result in the so called “hybrid vigour”, i.e., the tendency for hybrids to be superior to the parental genotypes ^44^. However, since hybrid vigour is related to heterozygosis, in our case we may speculate that strain-by-strain variability of regulons ^8^, as well as metabolic redundancy of *S. meliloti* genome ^4, 45, 46^(which could in some way mimic the presence of multiple alleles) could be the contributor to the increase in the observed symbiotic-related phenotype.

Summing up, these data highlight the potential of a large-scale genome manipulation approach to obtain highly effective, and cultivar specific, rhizobial strains. This provides a rational basis for the use of similar approaches in the development of elite bio-inoculants for use in precision agriculture ^12, 47^.

### Conclusions

The work presented here provides a proof-of-principle for the feasibility of using a large-scale genome manipulation approach that makes use of the species’ pangenome (i.e. the extended gene set present in a group of microbial strains belonging to the same species ^48^) to produce daughter strains with improved biotechnologically relevant (i.e., nitrogen fixing symbiosis) characteristics ^12^. In the current work, the large-scale genome manipulation was based on the transplantation of the primary symbiotic megaplasmid of a bacterial multipartite genome, a genome organization commonly found in the rhizobia. Although an entire replicon accounting for more than 20% of the total genome content was replaced with a homologous replicon of a closely related species, resulting in the gain of 482 new genes (in addition to numerous SNPs) and the loss of 354 genes, most of the core metabolic phenotypes appeared largely resilient to modification with this approach. However, others phenotypes, particularly complex (i.e. multigenic) phenotypes such as the symbiotic phenotypes, gave interesting features which support the validity of this approach to improve biotechnologically relevant properties.

## MATERIAL AND METHODS

### Microbiological and genetic methods

Strains and plasmids used in this study are described in Table 1. Conjugation between *E. coli* and *S. meliloti* were performed as described in ^49^. All growth media (LB, LBmc, TY, M9) and antibiotic concentrations were as previously described in^11, 45^.

### Cis-hybrid strain construction

First, a triparental mating between the wild type strain BL225C (the future donor), the helper strain *E. coli* MT616 (carrying pRK600 that has the RK2 *tra* genes) ^50^, and *E. coli* with the pTE3rctB vector (replicative plasmid overexpressing the *R. etli rctB* gene and carrying a tetracycline resistance marker) ^25^ was performed to create the BM848 (BL225C-rctB) strain. Secondly, a biparental mating between *S. meliloti* Rm3498 (∆pSymA) ^11^ and an *E.coli* S17-1 strain carrying the pMp7605 vector (carrying a gentamicin resistance marker) ^51^ was performed to generate the strain BM826. Lastly, the *cis*-hybrid strain BM806 was created through a biparental mating between the strain BM848 (BL225C-rctB) as the donor and the strain BM826 (ΔpSymA +pMp7605) as the acceptor. Selection for the *cis*-hybrid transconjugant strain (which had the pSymA replicon of the donor strain) was performed on M9 medium containing 1 mM MgSO_4_, 0.25 mM CaCl_2_, 0.001 mg/ml biotin, 42 µM CoCl_2_, 76 µM FeCl_2_, 10 mM trigonelline, streptomycin, and gentamycin. Streptomycin and gentamycin were used to select for the recipient strain, while the presence of trigonelline as the sole carbon source selected for the gain of pSINMEB01, as the trigonelline catabolic genes are located on pSymA/ pSINMEB01 ^52^.

### Validation of the transplanted strain

Pulsed-Field Gel Electrophoresis (PFGE) was performed to verify the successful uptake of pSymA via restriction digestion of genomic DNA with *Pme*I. The applied PFGE protocol was modified from Herschleb et. al 2007^53^ and Mavingui et al. 2002^54^, and a protocol from Sharon Long’s research group (Standford University, available at http://cmgm.stanford.edu/biology/long/files/protocols/Purification%20of%20S%20meliloti.pdf). *S. meliloti* cultures were grown to an OD600 of 1.0 in TY medium supplemented with suitable antibiotics and harvested by centrifugation (3000 *g*, 15 min, 4°C). All following steps were carried out either on ice or at 4°C. Sedimented cells were washed with TE buffer (10mM Tris-HCl, 1mM EDTA) supplemented with 0.1% (w/v) N-Lauroylsarcosine, and a second time with TE buffer. Washed cell pellets were then resuspended in TE buffer and mixed (1:1) with 1.6% (w/v) low-melt agarose (50°C), thereby resulting in a final concentration of ~8x10^8^ cells/ml. Two hundred µl of each suspension was casted into a moistened mold and gelatinized at 4°C. The resulting agar plugs were subsequently incubated at 37°C for 3 h in lysis buffer (6mM Tris-HCl, 1M NaCl, 100mM EDTA, 0.5% (w/v) Brij-58, 0.2% (w/v) Sodium deoxycholate, 0.5% (w/v) N-Lauroylsarcosine) supplemented with 1.5 mg/ml lysozyme (SERVA Electrophoresis GmbH, Germany). Treated agar plugs were then washed in H_2_O, followed by incubation at 50°C for 48 h in Proteinase K buffer (100mM EDTA, 10mM Tris-HCl, 1% (w/v) N-Lauroylsarcosine, 0.2% (w/v) Sodium deoxycholate, pH 8.0) supplemented with 1 mg/ml Proteinase K (AppliChem GmbH, Germany). Finally, agar plugs were sequentially washed in four steps, 1 h per wash. After incubation in washing buffer (10mM Tris-HCl, 50mM EDTA), plugs were washed in washing buffer supplemented with 1 mM Phenylmethylsulfonyl fluoride, then in washing buffer, and finally in 0.1x concentrated washing buffer.

For restriction digestion with *Pme*I (New England Biolabs, USA), the prepared agar plugs were incubated in 1 ml of restriction enzyme buffer (1x concentrated) for 1 h with gentle agitation at room temperature. Then, the plugs were transferred into 300 µl of fresh enzyme buffer supplemented with *PmeI* (50 units per 100 µl agar plug). Restriction digestions were incubated over night at 37°C. After overnight incubation, agar plugs were washed in 1x washing buffer for 1 h. For PFGE analysis, 1/8^th^ of each agar plug was used. PFGE was performed using the Rotaphor^®^ System 6.0 (Analytik Jena, Germany) following the manufactor’s instructions. Separation of DNA fragments was achieved using a 0.5% agarose gel (Pulse Field Certified Agarose, Bio-Rad, USA) and 0.5x TBE buffer (44.5 mM Tris-HCl, 44.5 mM boric acid, 1 mM EDTA). The following settings were applied: step 1 – 18 h, 130V-100V (logarithmic decrease), angle: 130°-110° (logarithmic decrease), interval: 50sec-175sec (logarithmic increase); step 2 – 18 h, 130V-80V (logarithmic decrease), angle: 110°, interval: 175sec-500sec (logarithmic increase); step 3 – 40 h: 80V-50V (logarithmic decrease), angle: 106°, interval: 500sec-2000sec (logarithmic increase). Buffer temperature was adjusted to 12°C.

For whole genome sequencing, a Nextera XT DNA library was constructed ^55^ and sequenced using the Illumina MiSeq platform which generated 2,504,130 paired-end reads. After trimming, assembly was performed with SPAdes 3.9.0 ^56^, which produced 399 contigs. Contigs were aligned against the genomes of *S. meliloti* 2011 and BL225C. The assembly has been deposited to the GenBank database under the BioProject ID PRJNA434498.

Finally, several PCR primer pairs for amplification of unique genes of Rm2011 and BL225C (Supplementary Table S3), selected based on a comparative genome analysis with Roary ^57^, were routinely used to ensure the correct identification of strains during all experiments.

### Growth curves

Growth curves were initiated by diluting overnight cultures to an OD_600_ of 0.1 in TY medium or in M9 medium supplemented with succinate as a carbon source. Incubation was performed in 150 µl volumes in a 96 well microtiter plate. The microplates were incubated without shaking’ at 30°C and growth was measured with a microplate reader (Tecan Infinite 200 PRO, Tecan, Switzerland).

### Growth with root exudates

The ability to colonize plant roots was tested using growth on root exudates as a metabolic proxy for colonization. Root exudates were produced from seedlings of *M. sativa* (cv. Maraviglia) as previously described in ^37^. Strains were grown on TY plates, following which a single colony was resuspended in 0.9% NaCl solution to a final OD600 of 0.5 (1×10^9^ CFU/ml). Then, each microplate well was inoculated with 75 μl of either M9 without a carbon source or a nitrogen-free M9 composition with succinate as a carbon source, 20 μl of root exudate, and 5 μl of the culture. The microplates were incubated without shaking’ at 30°C and the growth was measured on a microplate reader (Tecan Infinite 200 PRO, Tecan, Switzerland). At the end of the incubation period, aliquots from each well were diluted and viable titres of *S. meliloti* cells were estimated after incubation on TY plates at 30°C.

### Plant symbiotic assays

Symbiotic assays were performed in microcosm conditions in plastic pots containing a 1:1 mixture of sterile vermiculite and perlite, supplemented with 200 ml of Fahraeus N-free liquid plant growth medium ^58^. *S. meliloti* strains were grown in liquid TY medium at 30°C for 48 h. Cultures were then washed three times in 0.9% NaCl solution and resuspended to an OD_600_ of 1.0. Approximately 1×10^7^ cells were added to each pot, corresponding to ~ 4 × 10^4^ cells/cm^3^. Washed cell-suspensions were then directly spread over the roots of one-week old seedlings that were directly germinated in the pots, and grown in a growth chamber maintained at 26°C with a 16 h photoperiod (100 microeinstein/m^2^/s) for 5 weeks. Nodule counts were performed after the 5 weeks, then the shoots dried at 50°C for 7 days. The number of bacterial genome copies per nodule was determined with qPCR as previously reported ^40^. The alfalfa cultivars (*M. sativa*, *M. falcata*, *Medicago x varia*) used and their main features are reported in Supplemental Table S4.

### Biofilm assays

Strains were inoculated in 5 ml of TY and grown for 24 h with shaking. After growth, cultures were diluted to an OD600 of 0.02 in fresh TY medium and 100 µl of the diluted culture was inoculated into a microtiter plate. The plates were incubated at 30°C for 24 h, after which the OD600 was measured to determine the cell biomass. Each well was then stained with 20 μl of crystal violet solution for 10 minutes. The medium containing the planktonic cells was gently removed and the microtiter plate wells were washed three times with 200 μl of PBS (0.1 M, pH 7.4) buffer and allowed to dry for 15 min. The crystal violet in each well was then solubilized by adding 100 μl of 95% EtOH and incubating for 15 min at room temperature as described in ^59^. The plate was then read at 560 nm using a microtiter plate reader (Tecan Infinite 200 PRO, Tecan, Switzerland).

### Phenotype Microarray

Phenotype MicroArray™ experiments using Biolog plates PM1 (carbon sources), PM2A (carbon sources), and PM3 (nitrogen sources) were performed largely as described previously ^23^. All bacterial strains used in this study (parental and transplanted) are listed in Table 1. Data analysis was performed with DuctApe ^60^. Activity index (AV) values were calculated following subtraction of the blank well from the experimental wells. Growth with each compound was evaluated with AV values from 0 (no growth) to 4 (maximal growth), after elbow test calculation (Table S3 c;d).

### NMR metabolomics of the cell lysates and media

Overnight cultures were washed, resuspended, and diluted in 100 ml of fresh M9 medium (41 mM Na_2_HPO_4_, 22 mM KH_2_PO_4_, 8.6 mM NaCl, 18.7 mM NH_4_Cl, 4.1 µM biotin, 42 nM CoCl_2_, 1 mM MgSO_4_, 0.25 mM CaCl_2_, 38 µM FeCl_3_, 5 µM thiamine-HCl, 10 mM succinate) ^11^. For cell lysates, when cultures reached OD 1, 50 ml of each culture was pelleted by centrifuging for 25 minutes at 15000 *g*. For the media, 1 ml of supernatant of each culture was collected. For cell lysate analysis, each pellet was resuspended in 500 µL of PBS, and sonicated for 20 minutes with cycles of 1 second of activity and 9 seconds of rest (292.5 W, 13 mm tip), with contemporary cooling on ice. After cell lysis, the samples were centrifuged for 25 min at 4°C at 8000 *g*. For each strain, four independent experiments were performed. NMR samples were prepared in 5.00 mm NMR tubes (Bruker BioSpin) with 55 μL of a ^2^H_2_O solution containing 10 mM sodium trimethylsilyl[2,2,3,3-^2^H_4_] propionate (TMSP) and 500 μL of sample.

^1^H NMR spectra were acquired for both the cell lysates and the growth media. High reproducibility between samples was seen (Supplemental Figure S2), as expected based on previous studies with eukaryotic cells ^61^. NMR spectra were recorded using a Bruker 900 MHz spectrometer (Bruker BioSpin) equipped with a CP TCI ^1^H/^13^C/^15^N probe. Before measurement, samples were kept for 5 minutes inside the NMR probe head for temperature equilibration at 300 K. ^1^ H-NMR spectra were acquired with water peak suppression and a standard Carr–Purcell–Meiboom–Gill (CPMG) sequence (cpmgpr; Bruker BioSpin srl), using 192 or 256 scans (for cell lysates and growing media, respectively) over a spectral region of 18 kHz, 110 K points, an acquisition time of 3.07 s, and a relaxation delay of 4 s. This pulse sequence ^62^ was used to impose a T_2_ filter that allows selective observation of small molecular weight components in solutions containing macromolecules.

The raw data were multiplied by a 0.3 Hz exponential line broadening before applying Fourier transformation. Transformed spectra were automatically corrected for phase and baseline distortions and calibrated (chemical shift was referenced to the doublet of alanine at 1.48 ppm for cell lysates, and to the singlet of TMSP at 0.00 ppm for growth media) using TopSpin 3.5 (Bruker BioSpin srl). Multivariate and univariate analyses were performed on the obtained data using R software. For multivariate analysis, each spectrum in the region 10-0.2 ppm was segmented into 0.02 ppm chemical shift bins, and the corresponding spectral areas were integrated using the AMIX software (Bruker BioSpin). Binning is a mean to reduce the number of total variables and to compensate for small shifts in the signals, making the analyses more robust and reproducible. The area of each bin was normalized to the total spectral area, calculated with exclusion of the water region (4.50 – 5.15 ppm), in order to correct the data for possible differences in the cell count of each of the NMR samples.

Unsupervised Principal Component Analysis (PCA) was used to obtain a preliminary overview of the data (visualization in a reduced space, cluster detection, screening for outliers). Canonical analysis (CA) was used in combination with PCA to increase supervised data reduction and classification. Accuracy, specificity, and sensitivity were estimated according to standard definitions. The global accuracy for classification was assessed by means of a leave-one-out cross-validation scheme. The metabolites, whose peaks in the spectra were well defined and resolved, were assigned and their levels analyzed. The assignment procedure was performed using an internal NMR spectral library of pure organic compounds, public databases such as the *E. coli* Metabolome Database ^63^ storing reference NMR spectra of metabolites, and spiking NMR experiments ^64^. The relative concentrations of the various metabolites were calculated by integrating the corresponding signals in the spectra ^65^, using a home-made program for signal deconvolution. The nonparametric Wilcoxon-Mann-Whitney test was used for the determination of the meaningful metabolites: a p-value of 0.05 was considered statistically significant. The molecule 1,4-dioxane was used as a standard to perform the quantitative NMR analysis with the aim of obtaining the absolute concentrations (µM) of the analyzed metabolites.

NMR data were uploaded on the MetaboLights database (www.ebi.ac.uk/metabolights) with the accession number MTBLS576.

### Generation of the metabolic models

The manually curated iGD1575 reconstruction of *S. meliloti* Rm1021 ^19^ was modified to expand the composition of the biomass reaction through the inclusion of an additional 31 compounds, including vitamins, co-enzymes, and ions at trace concentrations as described elsewhere (Table S6) ^46^. Although iGD1575 is based on *S. meliloti* Rm1021, it is expected to accurately represent Rm2011 metabolism as these two strains are derived from the same field isolate (SU47) and have nearly identical gene contents ^20^; while there are numerous SNPs between the strains, SNPs are not considered during the process of metabolic reconstruction.

All other metabolic models were constructed using our recently published protocols for template-assisted metabolic reconstruction and assembly of hybrid bacterial models ^27^. Briefly, a draft metabolic reconstructions of *S. meliloti* BL225C was produced using the Kbase webserver (www.kbase.us) with gapfilling. The draft model was enhanced using the curated Rm1021 model as a template according to ^27^, using orthologous gene sets between BL225C and Rm1021 produced with InParanoid ^66^. Additionally, an appropriate *protein synthesis* reaction was manually added to the model. Finally, replicon transplantation between the BL225C model and the Rm1021 model was performed as described recently ^27^, making use of the InParanoid generated orthology data and the information contained within each model. All metabolic reconstructions used in this work are provided in Supplementary File S1 in COBRA format within a MATLAB MAT-file. The enhancement and transplantation pipeline is available at https://github.com/TVignolini/replicon-swap.

## Author contribution

A. Checcucci created the strains, performed microbiological analyses. G. diCenzo contributed in metabolic model creation and performed computation analyses on the metabolic modelling. V. Ghini, P. Turano, C. Luchinat performed NMR analyses and contributed in NMR spectra interpretation. V. Ghini contributed in preparing illustrations. A. Becker and J. Döhlemann contributed PFGE analysis and interpretation. T. Vignolini and M. Fondi contributed the first draft of metabolic model and preliminary computational simulations. G. diCenzo performed computational simulations. F. Decorosi and C. Viti contributed in Phenotype Microarray analysis and interpretation. A. Checcucci and C. Fagorzi contributed to *in vitro* symbiotic assays. M. Bazzicalupo and T. Finan provided data interpretation. A. Mengoni, M. Fondi, G. diCenzo, A. Checcucci conceived the work. A. Checcucci, A. Mengoni, V. Ghini, M. Fondi, G. diCenzo prepared the manuscript. All authors have read and approved the manuscript.

## Notes

Author declare no competing financial interest

## ACKNOWLEDGEMENTS

We are grateful to Dr. Carla Scotti (CREA-FLC, Lodi, Italy) for kindly providing seeds of *M. sativa* cultivars and to Gabriele Brazzini for assistance in symbiotic assays. NMR spectra were acquired and analyzed at CERM, Core Centre of Instruct-ERIC, an ESFRI Landmark, supported by national member subscriptions. This work was supported by the University of Florence, project “Dinamiche dell’evoluzione dei genomi batterici: l’evoluzione del genoma multipartito e la suddivisione in moduli funzionali”, call “PROGETTI STRATEGICI DI ATENEO ANNO 2014” to AM. AC was supported by a grant from Fondazione Buzzati-Traverso. GCD was supported by the Natural Sciences and Engineering Research Council of Canada (NSERC) through a PDF fellowship, VG was supported by Fondazione Umberto Veronesi. Work in the TMF lab is supported by NSERC. AB and JD were supported by the German Research Foundation (TRR 174).

## Supplemental Material

**Supplemental Table S1. List of primer used for the cis-hybrid strain creation**

**Supplemental Table S2. Differential utilization of carbon sources (a) and nitrogen sources (b) by cis-hybrid strain respect to the wild type strains.** The differential growth was evaluated with the activity index values calculated with DuctApe software (Galardini*, et al.*, 2014). Complete list of activity index values for all strains tested in carbon (c) and nitrogen (d) sources.

**Supplemental Table S3.Concentration (µM) of each identified metabolite (a) in the cell lysates and (b) in the media.**

**Supplemental Table S4. List of alfalfa cultivars used for symbiotic assay.**

**Supplemental Table S5. Symbiotic capabilities of the strains.** Values and relatives standard deviation of symbiotic performances profiles for Rm2011, BL225C and *cis*-hybrid strains in seven alfalfa cultivars.

**Supplemental Table S6. Biomass composition.**

**Supplemental File S1. Metabolic Model reconstruction in** COBRA format within a MATLAB MAT-file.

**Supplemental Figure S1. Pulse Field Gel Electrophoresis (PFGE)** performed on the cis-hybrid strains, the cured and the parental strains. The banding profile of the cis-hybrid strain is detailed with band size and replicon origin. The enzymatic digestion with PmeI was performed. Details: 0.5xTBE, 0.5% Agarose, M = PFGE marker *S. cerevisiae* (Biorad), Cell density: 8×10^8^/ml

**Supplemental Figure S2. NMR spectra of the endo (a) and exo (b) metabolic profiles.**

**Supplemental Figure S3. Boxplots of the interesting metabolites in cell lysates and media.**

**Supplementary Figure S4. Biofilm-to-biomass ratio** (OD560 / OD600) for the wild type, recipient, and cis-hybrid strains. The values are calculated on the mean values of 4 replicates. Error bars indicate standard deviation (calculated on the error propagation); (p <0.005, Tukey post-hoc contrasts).

**Supplemental Figure S5. Nodule colonization efficiency in *M.sativa* cv Maraviglia.** Nodule colonization (bacterial genome copies cells inside nodule, from qPCR, n=10); the experiment was performed in different biological replicate. Values indicate means and standard deviation (p <0.005, Tukey post-hoc contrasts).

